# Automated FRET Analysis for Enhanced Characterization of Protein-Protein Interactions

**DOI:** 10.1101/2025.09.20.674865

**Authors:** Ahmet Zübeyir Nursoy, Orkun Cevheroğlu, Çağdaş Devrim Son

## Abstract

Förster Resonance Energy Transfer (FRET) analysis is a powerful technique for studying protein-protein interactions; however, manual methods often introduce variability and user dependency. We present the SONLab FRET Analysis Tool, an open-source and automated software that integrates Cellpose for cell segmentation with standardized pipelines for bleed-through correction and FRET efficiency calculation. By minimizing human intervention, the tool improves reproducibility and comparability between experiments. The results demonstrate that the tool achieves FRET efficiencies comparable to those of manual methods but with reduced bias, enabling robust and high-throughput analysis of protein interactions.

**Research Highlights:**

- We developed an open-source pipeline using Cellpose for accurate, unbiased FRET analysis.
- It outperforms manual methods, reduces variability, supports high-throughput studies, and enables reproducible, user-friendly image analysis across labs.

**Graphical Abstract:** A visual summary of the SONLab FRET Tool, displaying the entire workflow—from imaging to the plotting of quantified data. The results provided in this article were obtained using this automated pipeline.

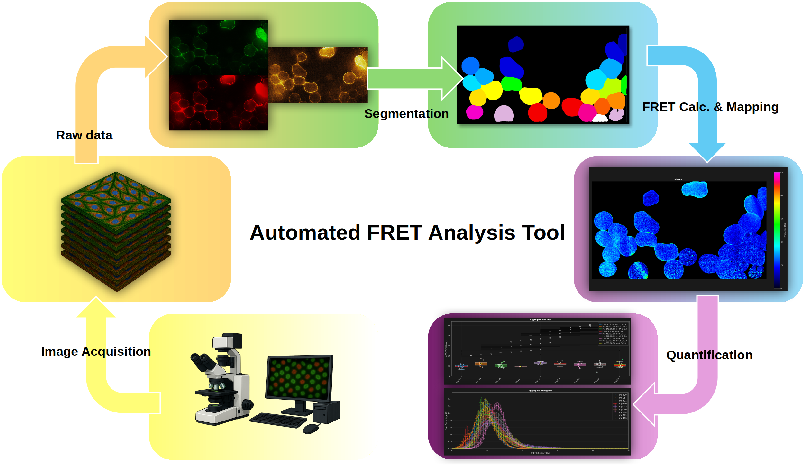

## Introduction

Förster Resonance Energy Transfer (FRET), also known as Fluorescence Resonance Energy Transfer, is a non-radiative dipole-dipole energy transfer from an excited fluorophore, mainly called the donor, to a compatible fluorophore, the acceptor. For efficient energy transfer, the donor’s emission spectrum has to overlap with acceptor’s excitation spectrum of the acceptor fluorophore, and the distance between fluorophores should typically be less than 10 nm. These properties make FRET a widely used technique for detecting molecular interactions and measuring nanometer-scale distances between interacting bodies (Jares-Erijman and Jovin, 2003; Sun *et al*., 2013).

The most commonly used intensity-based FRET technique is called 3-filter (cube) FRET (3F-FRET), also known as sensitized emission FRET, which is designed to quantify the amount of acceptor emission resulting from donor to acceptor energy transfer (Hochreiter *et al*., 2019). 3F-FRET involves three individual combinations of excitation wavelengths and emission filters:i) a donor detection channel (donor-specific excitation with a donor-specific emission filter), an acceptor channel (acceptor-specific excitation with ii) an acceptor-specific emission filter), iii) a raw FRET channel (donor-specific excitation with an acceptor-specific emission filter).

Among the earliest and most widely adopted tools for analyzing 3F-FRET data is PixFRET, an ImageJ plug-in that succeeded earlier software such as RiFRET (Roszik *et al*., 2009), PFRET (Wallrabe *et al*., 2015). PixFRET was designed for the pixel-by-pixel analysis of sensitized-emission FRET, aiming to provide quantitative and spatially resolved information about protein-protein interactions by calculating and displaying normalized FRET images. A key feature of PixFRET is the correction for spectral bleed-throughs (SBTs) and normalization for varying donor and acceptor expression levels. PixFRET addresses SBT variations by allowing users to model SBT ratios as a function of fluorophore intensity (through linear or exponential fitting) or by applying constant values derived from image stacks of cells expressing only the donor or acceptor fluorophore. After background subtraction, the raw FRET signal is corrected using these calculated SBT parameters. PixFRET then provides several normalization options for the corrected FRET signal, including division by donor intensity, acceptor intensity, the product of donor and acceptor intensities, the square root of their product, or the calculation of FRET efficiency 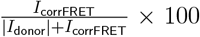. To ensure accuracy, FRET values are computed only for pixels where the signals in each channel and the product of donor and acceptor signals exceed specific background-derived thresholds, with non-accepted pixels visually indicated in blue. An optional Gaussian blur filter can be applied to smoothen the images, which helps in reducing noise and preventing aberrant FRET values, thereby improving the quality of normalized FRET images (Feige *et al*., 2005).

While FRET technique can provide robust quantitative information about protein interactions with the properly normalized raw data, one of the greatest challenges arises from variations in donor and acceptor expression levels in transfected cells. To address this, Hochreiter et al. introduced DFRET Hochreiter *et al*. (2019) providing an alternative normalization approach,conceptually similar to PixFRET (Feige *et al*., 2005). DFRET provides more robust results when compared to Youvan *et al*. (1997), Gordon *et al*. (1998), Xia and Liu (2001)’s earlier methods.

The DFRET method offers a significant advancement over traditional FRET normalization techniques such as NFRET and FRETN. It provides quantitative, spatially resolved analysis of protein-protein interactions in living cells, particularly taking into account interaction stoichiometries and relative affinities. In contrast to classical normalization methods, which mathematical simulations and live-cell data demonstrate often fail to produce a saturation plateau at high acceptor concentrations -and may even decline or yield inconsistent results-DFRET accurately reflects the progressive saturation of donor molecules following the law of mass action. Methodologically, DFRET uses a 3-filter FRET approach with specific correction factors (C1 and C2) that empirically relate the signal intensity of the FRET channel to those of the donor and acceptor channels. These factors are determined using a tandem fusion constructs with validated transfer efficiency, often by acceptor photobleaching, allow calculation of apparent FRET efficiencies and relative donor/acceptor concentrations for each individual cell. This normalization enables the plotting of FRET saturation curves, which can be fitted to a mathematical model to extract critical biophysical properties such as the interaction stoichiometry (z), maximum FRET efficiency (FRETmax) at complete donor saturation, and a relative binding affinity 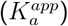. The ability to obtain such robust, independent, and quantitative measures across a broad range of donor/acceptor ratios, particularly when combined with large datasets from methods like flow cytometry, overcomes the key limitations of traditional FRET measures, making DFRET suitable for advanced evaluation of protein interactions that were previously unattainable (Hochreiter *et al*., 2019). While DFRET itself is a powerful and high-throughput method once calibrated, the demanding nature of its photobleaching-based calibration step limits its universal accessibility. That is why it was not implemented into the tool, but can be easily implemented thanks to its modular nature.

Rather than going into the details of the efficiency and robustness of these derived formulas, which is abundantly discussed in the corresponding literature, we would like highlight the dependency on the analyzer that prepares the data up to the point where aforementioned formulas come into the picture. We observed that analyzer dependency and “eyeballing” problems cause inaccurate and incomparable results across different experiments and instruments since the changing parameters do not only come from experimental or instrument parameter differences but also unmeasurable and untraceable factors that are introduced by the analyzer.

In many conventional software tools like the PixFRET ImageJ plug-in (Feige *et al*., 2005), users manually draw regions of interest (ROIs) on images. This process is subjective, leading to variability across users—or even within the same user at different times, introducing variability in the data. Eventually, this results in the analyzer dependency, which brings about changing inclusion criteria for data points, which are mostly cells or cellular compartments. No two researchers, even within the same group, are likely to select the same regions. Additionally, defining ROIs is laborious and time consuming, limiting the number of data points, while whole frame analysis often lacks sufficient spatial resolution.

To overcome these issues, we propose a “comprehensive (**and automated**) all-in-one software solution” designed to standardize **and automate** the FRET analysis pipeline, minimize user dependency and improve data comparability and robustness of results.

Given all these, we also acknowledge the persistent limitations of FRET, such as detector and instrument specific inconsistencies, qualitative and instrument-dependent outputs, issues like the orientation factor (*κ*^2^). However, by reducing human-induced variability and making analysis steps transparent and traceable, our tool enables reproducible and reliable cross-comparison of FRET datasets across experiments and laboratories.

## Methods

The SONLab FRET Analysis Tool is designed to aim a comprehensive, all-in-one software solution to standardize and automate the complete FRET analysis pipeline, thereby minimizing user dependence and improving data comparability and robustness.

### Integration of Cellpose for Automated Cell Segmentation

One of the fundamental aspects of the tool is the seamless integration with Cellpose (Stringer *et al*., 2021), a state-of-the-art deep learning-based cell segmentation method developed by Stringer and colleagues. Cellpose leverages neural networks for accurate and precise detection of cell boundaries, providing an automated and configurable process that was traditionally done manually via “eyeballing.” Cellpose significantly reduces user-induced variability introduced by manual ROI drawing, and enables “cell-specific analysis” and “batch processing for high-throughput analysis “with “consistent segmentation parameters” across multiple images.

The software allows users to choose the model that Cellpose offers, minimum cell diameters, flow threshold, and cell probability threshold. Following segmentation, post-processing is applied to remove small objects or cells that cannot be included as data points and when appropriate, to delineate cellular membranes based on the localization of the molecules of interest.

All User Interface tabs are designed with PyQt5 (Riverbank-Computing), and plots are generated using Matplotlib (Hunter, 2007). Post-processing steps are performed with Open-CV (Bradski, 2000) and Scikit-Image (Van der Walt *et al*., 2014), while images and data are handled by czifile and tifffile (Gohlke, 2024), depending on the input image format, alongside core numerical operations implemented with Numpy (Harris *et al*., 2020).

In cell segmentation tab, users can adjust Cellpose and post-processing parameters. Since the segmentation might detect cells that do fit the user’s criteria of inclusion, ROIs that do not meet inclusion criteria can be removed or some regions can be added manually. After segmentation is satisfactory, based on the provided visualization of the original and the segmented mask images, results are saved as a TIF file with an additional first frame to the stacked original frames into the distinct directory of each input with proper naming and might be transferred to the analysis tab directly. After parameter optimization for a given dataset, batch segmentation can be applied to accelerate large-scale analyses. The tool natively supports CZI files, the output format of Zeiss microscopes used in this study, by converting them to TIF format without data loss. More generally, the tool is not only limited with the file type, any microscopy image format can be analyzed once converted to TIF. The entire pipeline of the segmentation tab is illustrated in Figure 1.

**Figure 1.**
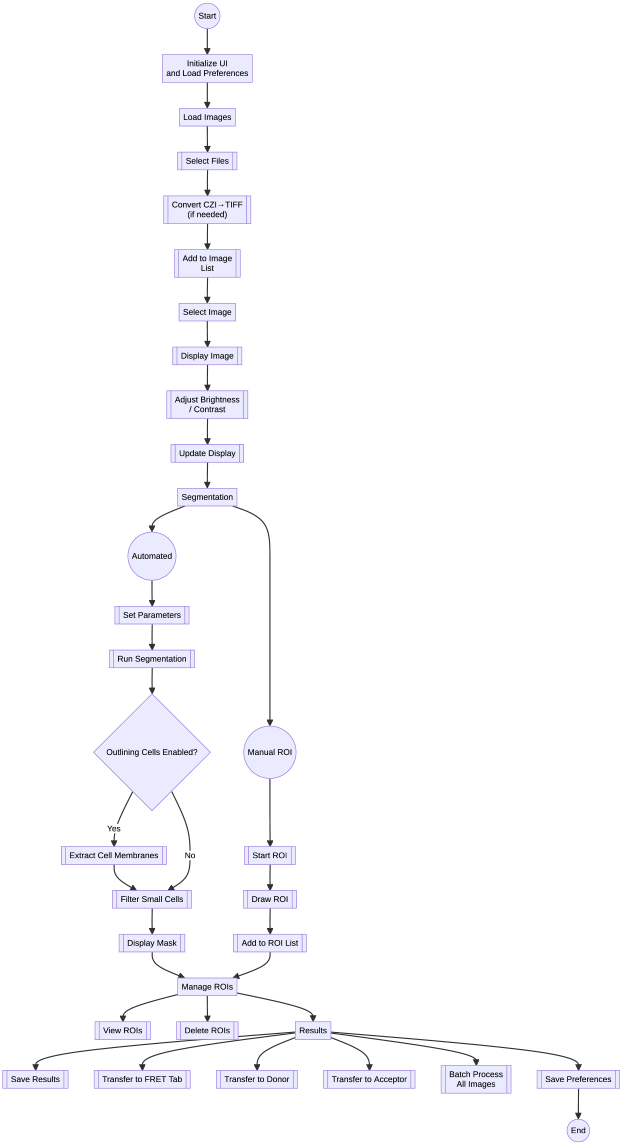
General pipeline of the Cellpose integrated segmentation tab.

In the analysis tab, output of the segmentation tab is required in the form of a four-frame TIF file containing: i) labels, ii) FRET channel, iii) Donor channel, and iv) Acceptor channel.

### Bleed-Through (BT) Calculation

This tab provides a PixFRET (Feige *et al*., 2005) like implementation for bleed-through calculation, in which equations are fitted separately for donor and acceptor BTs into the FRET channel in two separate sub-tabs. Two parameters are calculated for each pixel in donor-only and acceptor-only samples: *S*_1_ for donor-only and *S*_2_ for acceptor-only:

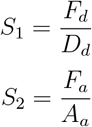

For donor and acceptor, these values are plotted against corresponding donor and acceptor intensities. The software then fits three different models; constant, linear, and exponential, to the data. Users can visually inspect the fits and select the model that best represents their dataset for each case. An example fit on the plot is shown in Figure 2

**Figure 2.**
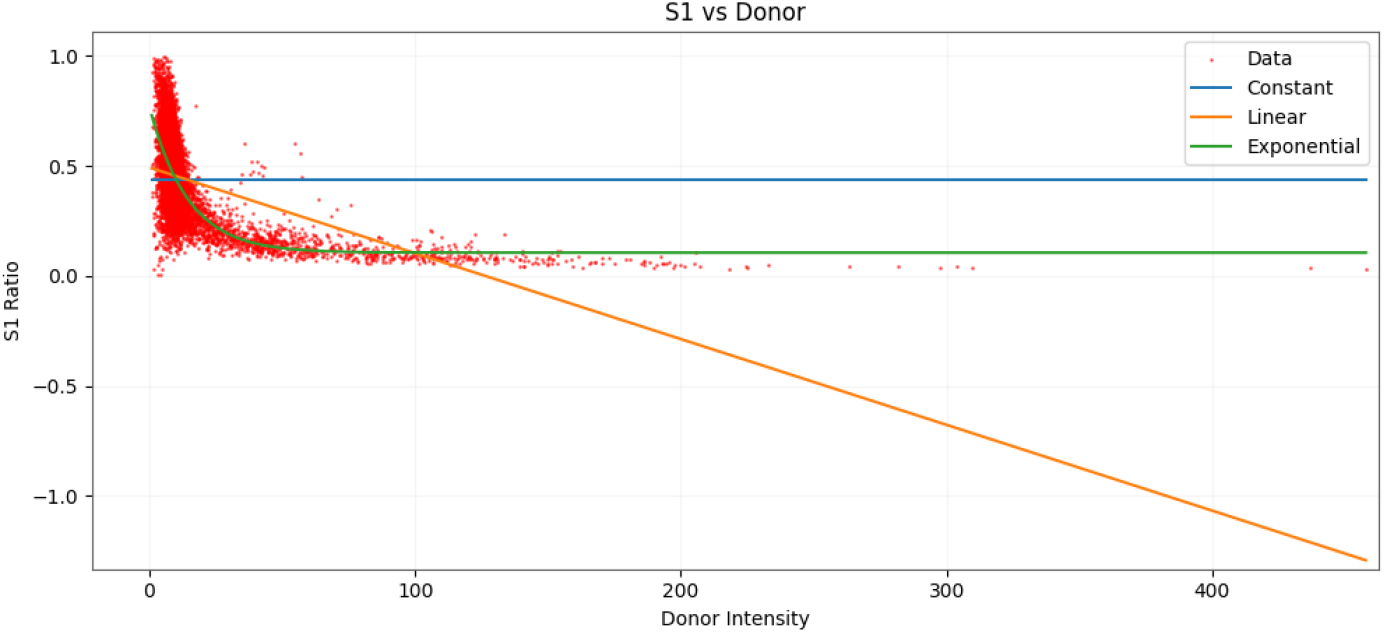
An example scatter plot of S1 vs Donor intensity. This plot provides information about how BT is changing with the changing donor intensity. The fit equations are provided, constant, linear, and exponential. For better fittings, data points to be fitted can be adjusted by dynamically defining the max and min points of the data to be included.

For curve fitting, a specific area on the plot can be selected by the interactive border determination feature instead of traditional zooming or panning, which is untraceable, to obtain more accurate fitting equations. The overall workflow of the BT tab is illustrated in Figure 3. To improve model reliability, multiple images as can be added and analyzed within the same fitting process.

**Figure 3.**
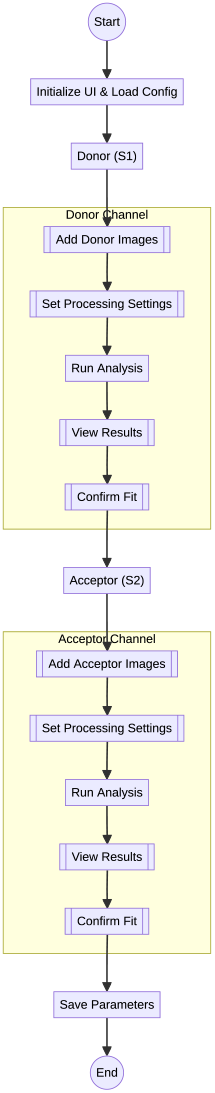
Flow of the bleed through calculation tab’s pipeline.

### FRET Analysis and Visualization

The FRET analysis tab is the heart of the tool. It consists of two sub-tabs: one provides colored efficiency maps based on user-selected parameters, plotting efficiency images for any selected file from the image list; the other provides summary statistics either per cell or for all images grouped by experimental condition (e.g., Positive Ctl, Negative Ctl etc.). Statistical tests (e.g., ANOVA, t-test) are done by using Scipy library (Virtanen *et al*., 2020), and Gaussian mixture modelling (GMM) from Scikit-Learn (Pedregosa *et al*., 2011) is applied to assess the distribution of FRET efficiencies within a single cell. The workflow of the FRET tab is illustrated in Figure 4.

**Figure 4.**
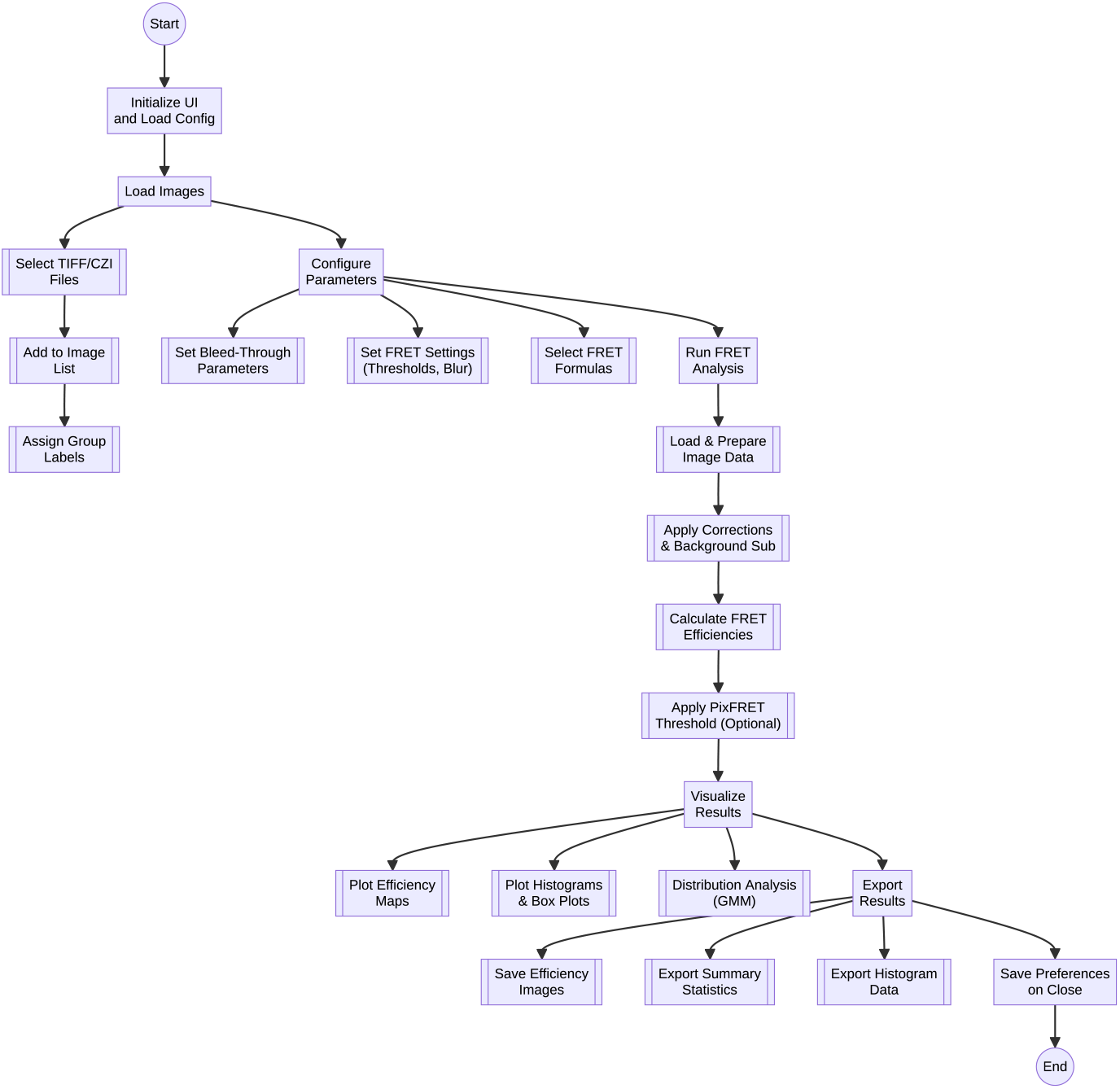
FRET Analysis Tab’s workflow. Almost all visualizations and statistics are exportable for any downstream use.

Background (BG) signal is determined by a moving kernel with a user defined size to find the minimum mean value using an uniform filter and subtracted from the pixel values for each frame of the input image. This eliminates analyzer-dependent BG signal determination and ensures consistent subtraction across different users analyzing the same type of image sets. For direct comparison with the widely used PixFRET method, its formula and thresholding logic are also implemented in the tool.

To calculate normalized FRET efficiency, five distinct formulas are implemented, each based on the corrected FRET signal, *F*^*c*^. The variables are defined as follows:

- *F*^*r*^: Raw FRET channel intensity (donor excitation, acceptor emission).
- *D*^*r*^: Raw donor channel intensity (donor excitation, donor emission).
- *A*^*r*^: Raw acceptor channel intensity (acceptor excitation, acceptor emission).
- *D*^*c*^: Corrected donor channel intensity.
- *A*^*c*^: Corrected acceptor channel intensity.
- *F*^*c*^: Corrected FRET channel intensity, accounting for spectral bleed-through.
- *F*_*d*_: FRET channel intensity from donor-only samples.
- *D*_*d*_: Donor channel intensity from donor-only samples.
- *F*_*a*_: FRET channel intensity from acceptor-only samples.
- *A*_*a*_: Acceptor channel intensity from acceptor-only samples.
- *G*: A constant used in normalization, specific to the experimental setup (Gordon *et al*., 1998).

The corrected FRET signal, *F*^*c*^, is computed by subtracting spectral bleed-through contributions from the donor and acceptor, assuming negligible bleed-through between donor and acceptor channels (i.e., raw intensities equal corrected intensities: *D*^*r*^ = *D*^*c*^, *A*^*r*^ = *A*^*c*^). The correction formula is:

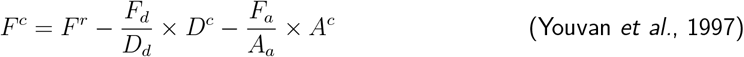

Using *F*^*c*^, the normalized FRET efficiency is calculated using one of the following five formulas, each representing a different normalization approach. The results from these formulas can be visualized as histograms or box plots in software’s analysis tab, enabling side-by-side comparison and statistical testing (e.g., ANOVA, t-test):

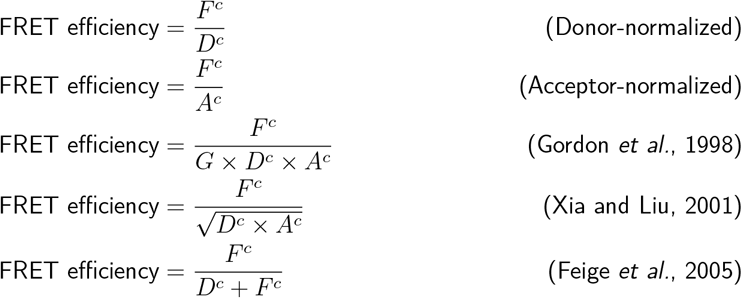

Each formula normalizes the corrected FRET signal differently:

- **Donor-normalized**: Normalizes by donor intensity, emphasizing donor contribution.
- **Acceptor-normalized**: Normalizes by acceptor intensity, emphasizing acceptor contribution.
- **Gordon et al**.: Accounts for both donor and acceptor intensities with a scaling factor *G*.
- **Xia et al**.: Uses the geometric mean of donor and acceptor intensities for normalization.
- **Feige et al**.: Represents FRET efficiency as a fraction of the total donor signal posttransfer.

After calculating FRET using the given formulas above, values below the lower threshold or above the upper threshold are excluded from mean and statistical analyses. Lower and upper thresholds are generally determined by the fluorescent pair used. The upper threshold is particularly useful for excluding implausibly high efficiency values that exceed the theoretical maximum.

In addition to classical box plots for each image or each group with statistical comparisons, t-test for two groups and ANOVA for more than two, aggregate histograms are provided for both individual images and all images in a group by calculating means of cells that they contain. These histograms summarize the distribution of FRET efficiencies across cells, expressed as percentages of pixel ratios. This helps assess the uniformity of interactions within cells. To ensure smooth visualization, efficiencies are divided into 256 bins, providing sufficient resolution for percent FRET efficiency in 16-bits of images.

## Results

To evaluate the effectiveness of the developed tool, two representative groups commonly used in the FRET analysis were analyzed as a showcase: a positive control, Gap43 (mCherry-Linker-mEGFP), and negative (soluble mCherry and mEGFP). An detailed summary of the analysis is presented in Figure 5.

**Figure 5.**
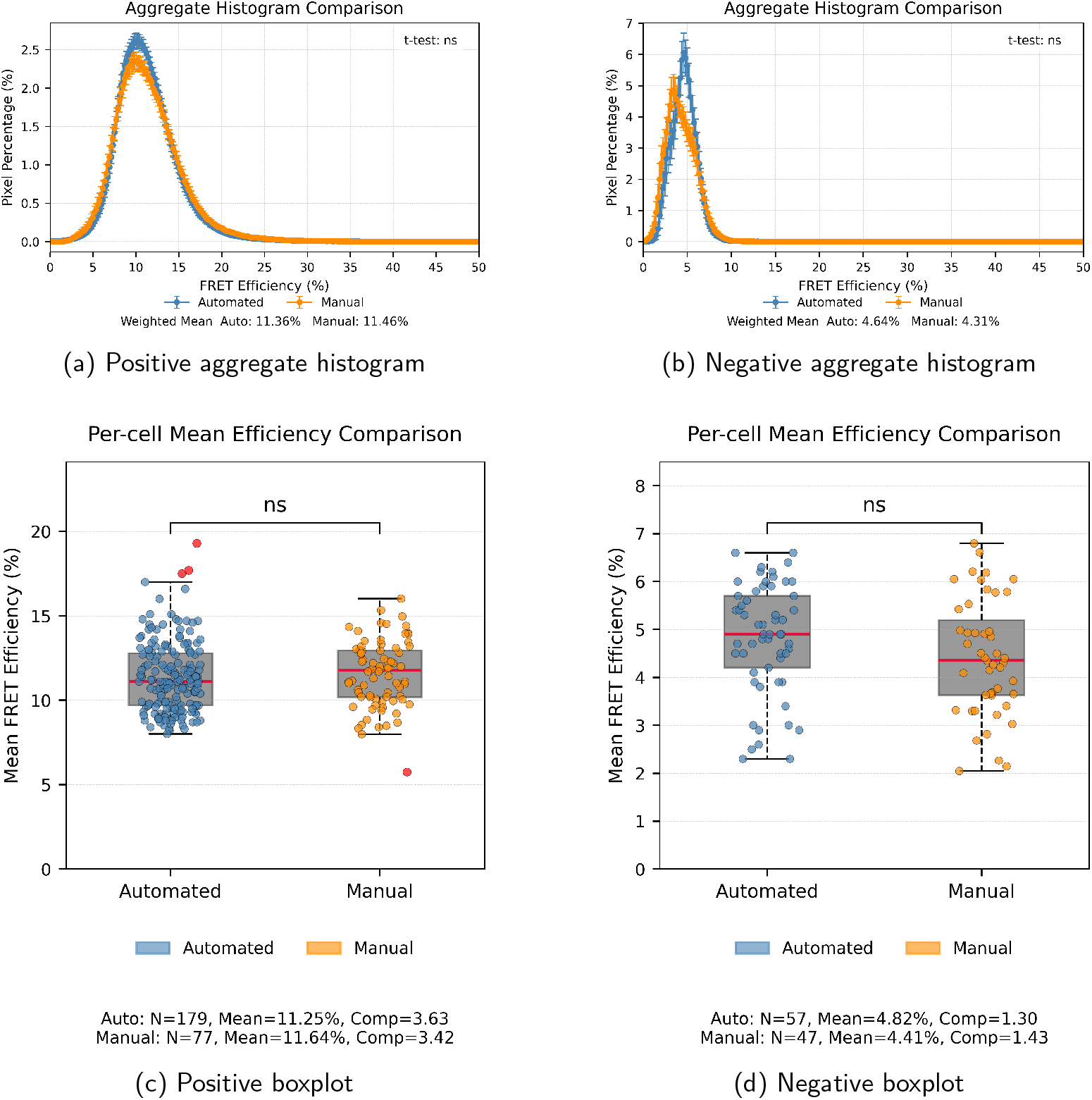
Comparison of aggregate histograms and boxplots for positive and negative classes.

For the same image set, manual analysis was performed by user-drawn ROIs and calculating FRET efficiencies using ImageJ with the PixFRET plug-in. In this case, number of cell (*N*) included was 77 for the positive control samples and 47 for the negative control. The average FRET efficiency was calculated as 11.64% and 4.41% for positive and negative, respectively. To determine the deviation of the cells from the mean, variance was measured as variance 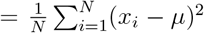 where *x*_*i*_ is an inlier and *µ* is the mean. The variance was 1.43 for the negative control set and 3.42 for the positive control set.

Using the same image sets and the formulas-but without the PixFRET correction factor implementation-the automated software identified 179 cells in positive controls group and 57 in negative controls group. The resulting average FRET efficiencies were calculated as 11.25% and 4.82% with compactness of 3.63 and 1.30 for the positive and negative sets, respectively.

To assess consistency between automated and manual analysis done by ImageJ and PixFRET plug-in, t-tests were performed for both positive and negative sample groups. No statistically significant differences were observed in either the positive (p = 0.1332) or negative (p = 0.0779) sets.

**Table 1:** Comparison of FRET Efficiency Analysis: Manual vs. Automated Methods.

| Parameter | Positive Control | Negative Control |
| --- | --- | --- |
| *Manual Analysis (ImageJ with PixFRET)* |  |  |
| Number of Cells ($N$) | 77 | 47 |
| Average FRET Efficiency (%) | 11.64 | 4.41 |
| Compactness | 3.42 | 1.43 |
| *Automated Analysis (Software without PixFRET Correction)* |  |  |
| Number of Cells ($N$) | 179 | 57 |
| Average FRET Efficiency (%) | 11.25 | 4.82 |
| Compactness | 3.63 | 1.30 |
| *Statistical Analysis* |  |  |
| t-test p-value | 0.1332 | 0.0779 |

## Discussion

The most evident limitation of the manual analysis lies in the subjective selection of the cells, the variation of them and particularly in positive sets. Some researchers define ROIs directly on the analyzed FRET efficiency frame where interactions are already visible. This approach introduces personal bias (“eye-balling”) as experienced users might tend to select cells that appear to fall within the expected efficiency range, especially when analyzing positive and negative controls. Consequently, greater variability was observed in automated analysis compared with manual analysis for positive controls.

In negative sets, manual and automated analyses converge to highly similar results. This showed that nearly all visible cells were included 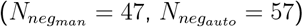 with minimaldiscrepancy, as expected for negative controls. Importantly, the robustness and effectiveness of the software become more evident as the dataset size increases. While the tool performs well even with small sample numbers; such cases require careful cell segmentation and batch segmentation is best avoided. With sufficiently large datasets, however, batch segmentation reliably manages to extract the characteristics of the interactions of interest regardless.

For both negative and positive samples, the software tool provided similar results that align closely with manual analysis in terms of interaction levels. Given that, we emphasize that the software’s results are more likely to reflect the true biology, as they are free from cherry-picking of data or inconsistent, untraceable parameter determination. Once the parameters are optimized for a specific instrument and experimental setup, cross-experiment comparisons become feasible. Experiment groups can then be interpreted by comparing to positive and negative controls without the introduction of the personal bias, which cannot be uniformly applied across conditions.

Notably, in the positive control, the software identified substantially larger number of cells 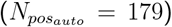 compared with the manual selection 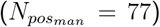 . The software stillm managed to extract the overall characteristics of the data set, while eliminating the considerable manual effort required from the analyzer, and freeing their time for image acquisition or segmentation optimization.

The slight numerical differences between manual and automated calculation by the software can be attributed to differences in the number of included cells, different background signal determination, different bleed-through fit or coefficients, and correction factor (CF) logic used by PixFRET. Although PixFRET’s CF implementation is also available in our software for direct comparison, we did not use it here, as Cellpose-based segmentation adequately fulfils the intended purpose the usage of CF.

The results presented here serve as a proof of concept and as a validation of correct methodology rather than demonstration of all the features of the developed software. The software has been made fully accessible to the community, and we encourage researchers to use, evaluate, and contribute to its ongoing development.

## Conclusion

The purpose of developing this software is not to introduce a new method for FRET analysis, but rather to integrate existing methods into a seamless pipeline while minimizing the influence of human factors-one of the most critical issues affecting reproducibility. The all-in-one software, which is fully open source and available via GitHub, enables researchers to reduce analysis time from weeks to hours. Even researchers with no coding knowledge can make a use the software easily. Importantly, the tool is designed to be user-friendly, allowing even those without coding experience to use it effectively. We believe this platform will serve as a backbone for future improvements and enhancements contributed by the scientific community. Moreover, it provides a foundation for incorporating deep learning (DL) and machine learning (ML) models into FRET analysis workflows, further demonstrating their potential for robust and efficient analysis.

Currently, there is a notable lack of tools that combine deep learning with traditional three-channel, image-based FRET analysis. This gap represents a clear opportunity for innovation. By leveraging established platforms like ImageJ and its plugins such as RiFRET or PixFRET, it is possible to develop hybrid solutions that combine the reliability of proven methods with the power of automation and modern computational approaches. Such developments would directly address the growing demand for more reliable, user-friendly tools capable of handling increasingly complex microscopy datasets, thereby bridging the gap between cutting-edge machine learning and proven FRET analysis methodologies.

## Data Availability

The software developed in this study is openly available at https://github.com/sonlab-metu/SONLab-FRET-Tool. Example datasets and analysis scripts used for validation are available from the corresponding author upon reasonable request.

## Ethics Approval and Consent to Participate

Not applicable. This study did not involve human participants or animal experiments.

## Acknowledgement

This study was supported within the scope of the Middle East Technical University Multi-disciplinary Research Project Programme with project number of ÇDAP-108-2025-11559 and TUBITAK research grant 118Z694. We are also thankful to Asst. Prof. Aybar Can ACAR and Prof. Ahmet Bugra KOKU for their insights and suggestions.

## References

Bradski, G. (2000). The OpenCV Library. Dr. Dobb’s Journal of Software Tools.

Feige, J. N., Sage, D., Wahli, W., Desvergne, B., and Gelman, L. (2005). Pixfret, an imagej plug-in for fret calculation that can accommodate variations in spectral bleed-throughs. Microscopy research and technique, 68(1):51–58.

Gohlke, C. (2024). cgohlke/tifffile: v2024.5.3.

Gordon, G. W., Berry, G., Liang, X. H., Levine, B., and Herman, B. (1998). Quantitative fluorescence resonance energy transfer measurements using fluorescence microscopy. Biophysical journal, 74(5):2702–2713.

Harris, C. R., Millman, K. J., van der Walt, S. J., Gommers, R., Virtanen, P., Cournapeau, D., Wieser, E., Taylor, J., Berg, S., Smith, N. J., Kern, R., Picus, M., Hoyer, S., van Kerkwijk, M. H., Brett, M., Haldane, A., del Río, J. F., Wiebe, M., Peterson, P., Gérard-Marchant, P., Sheppard, K., Reddy, T., Weckesser, W., Abbasi, H., Gohlke, C., and Oliphant, T. E. (2020). Array programming with NumPy. Nature, 585(7825):357–362.

Hochreiter, B., Kunze, M., Moser, B., and Schmid, J. A. (2019). Advanced fret normalization allows quantitative analysis of protein interactions including stoichiometries and relative affinities in living cells. Scientific reports, 9(1):8233.

Hunter, J. D. (2007). Matplotlib: A 2d graphics environment. Computing in Science & Engineering, 9(3):90–95.

Jares-Erijman, E. A. and Jovin, T. M. (2003). Fret imaging. Nature biotechnology, 21(11):1387–1395.

Pedregosa, F., Varoquaux, G., Gramfort, A., Michel, V., Thirion, B., Grisel, O., Blondel, M., Prettenhofer, P., Weiss, R., Dubourg, V., et al. (2011). Scikit-learn: Machine learning in python. Journal of machine learning research, 12(Oct):2825–2830.

Riverbank-Computing. Pyqt5. https://www.riverbankcomputing.com/software/pyqt/. Accessed: 2025-07-21.

Roszik, J., Lisboa, D., Szöllősi, J., and Vereb, G. (2009). Evaluation of intensity-based ratiometric fret in image cytometry—approaches and a software solution. Cytometry Part A, 75(9):761–767.

Stringer, C., Wang, T., Michaelos, M., and Pachitariu, M. (2021). Cellpose: a generalist algorithm for cellular segmentation. Nature methods, 18(1):100–106.

Sun, Y., Rombola, C., Jyothikumar, V., and Periasamy, A. (2013). Förster resonance energy transfer microscopy and spectroscopy for localizing protein–protein interactions in living cells. Cytometry Part A, 83(9):780–793.

Van der Walt, S., Schönberger, J. L., Nunez-Iglesias, J., Boulogne, F., Warner, J. D., Yager, N., Gouillart, E., and Yu, T. (2014). scikit-image: image processing in python. PeerJ, 2:e453.

Virtanen, P., Gommers, R., Oliphant, T. E., Haberland, M., Reddy, T., Cournapeau, D., Burovski, E., Peterson, P., Weckesser, W., Bright, J., van der Walt, S. J., Brett, M., Wilson, J., Millman, K. J., Mayorov, N., Nelson, A. R. J., Jones, E., Kern, R., Larson, E., Carey, C. J., Polat, İ., Feng, Y., Moore, E. W., VanderPlas, J., Laxalde, D., Perktold, J., Cimrman, R., Henriksen, I., Quintero, E. A., Harris, C. R., Archibald, A. M., Ribeiro, A. H., Pedregosa, F., van Mulbregt, P., and SciPy 1.0 Contributors (2020). SciPy 1.0: Fundamental Algorithms for Scientific Computing in Python. Nature Methods, 17:261– 272.

Wallrabe, H., Sun, Y., Fang, X., Periasamy, A., and Bloom, G. S. (2015). Three-color confocal förster (or fluorescence) resonance energy transfer microscopy: Quantitative analysis of protein interactions in the nucleation of actin filaments in live cells. Cytometry Part A, 87(6):580–588.

Xia, Z. and Liu, Y. (2001). Reliable and global measurement of fluorescence resonance energy transfer using fluorescence microscopes. Biophysical journal, 81(4):2395–2402.

Youvan, D. C., Coleman, W. J., Silva, C. M., Petersen, J., Bylina, E. J., and Yang, M. M. (1997). Fluorescence imaging micro-spectrophotometer (fims). Biotechnology et alia, 1:1– 16.

